# Quantification of domain-specific intrinsic capacity using mortality data

**DOI:** 10.64898/2026.04.06.714260

**Authors:** Matías Fuentealba, Ting Zhai, Saleem A Al Dajani, Vadim N Gladyshev, Michael P Snyder, David Furman

**Author notes:** Corresponding author: David Furman, PhD.

## Abstract

Functional health is centered on five domains of Intrinsic Capacity (IC): locomotion, cognition, vitality, psychological and sensory capacity. Therefore, measuring IC at the domain-specific level is the cornerstone for developing preventive interventions to help individuals preserve their independence. In this study, we used 63 clinical features from the UK Biobank to develop ‘IC age’, an 18-year mortality risk estimator that approximates an individual’s biological age associated with the decline of each IC domain. By establishing proteomic surrogates of IC age, we find immune system activation across domains and provide a proteomic framework that may facilitate scalable monitoring of functional health decline.

## Main

The global demographic shifts in the aging population require a fundamental transformation in the healthcare framework, shifting from reactive and disease-centric models towards proactive strategies that preserve function as we age. Central to this transformation is the concept of Intrinsic Capacity (IC), introduced by the World Health Organization (WHO) in 2015 and defined as the composite of an individual’s physical and mental capabilities^1,2^. IC is composed of five domains, including locomotor capacity (physical movement); sensory capacity (such as vision and hearing); vitality (energy and strength); cognition; and psychological wellbeing, all of which are modulated by environment and lifestyle to determine functional abilities including meeting basic needs; learning, making decisions; being mobile; building relationships; and contributing to society^3^. In recent years, studies have validated IC as a robust predictor of future clinical outcomes, including multimorbidity^4–6^, quality of life^7,8^, age-related diseases such as dementia^9^, cardiovascular^10^ and respiratory^11^ diseases, as well as hospitalization^12,13^ and all-cause mortality^12,14–16^.

Measuring IC currently relies on a combination of clinical assessments and emerging molecular tools. For example, the Mini-Mental State Examination (MMSE) remains the most frequent test to measure cognition^17^. Clinical assessments of vitality generally use nutritional^4,18,19^, metabolic^6^ and energy-related indicators^8,20^. For locomotion, practitioners typically use physical performance tests^18,21,22^. Similarly, depression scales evaluate psychological capacity^23,24^, whereas clinical tools^4,25^ and self-reported hearing and visual levels^6,20^ quantify sensory capacity. Beyond these clinical assessments, emerging molecular tools such as the IC clock^26^, use blood epigenetic data to predict intrinsic capacity based on the methylation levels of 91 CpGs.

Similar to biological aging, intrinsic capacity is not uniform, as individuals experience functional decline in different domains of IC^27^ as well as in organs^28,29^. Thus, monitoring specific IC domains allows the identification of individuals at risk prior to the manifestation of diseases such as cognitive decline and Alzheimer’s disease, or a decrease in locomotion capacity and sarcopenia or osteoporosis, while also providing a window for targeted treatment based on the associated decline in specific domains. For example, if risk is driven primarily by the locomotion domain, clinicians can tailor interventions toward physical therapy or nutritional supplementation rather than a generic wellness plan.

This study leverages clinical features describing IC to develop domain-specific mortality risk estimators designed to represent the biological age of each IC domain (‘IC age’). We performed association analyses to evaluate these metrics against disease incidence, multi-morbidity, medication and lifestyle behaviors such as diet and sleep to identify distinct stressors influencing each domain. To improve the mechanistic understanding and clinical applicability, we identified proteomic surrogates of domain-specific IC age predictors, enabling IC decline assessment from a blood sample and making the measurement scalable for population-level public health monitoring.

We integrated 63 IC domain-specific features from the UK Biobank into Cox Proportional Hazard models to develop mortality risk predictors with follow-up time up to 18.3 years for each of the five domains of intrinsic capacity **(Sup Table. 1-2)**. To enhance interpretability, we transformed the relative mortality risk estimates into age values by matching their distribution with chronological age, resulting in the IC age **(Fig. 1a)**. As expected, IC age displays high correlations with chronological age across all domains. We observed the strongest correlation in the sensory domain (R = 0.96, C-index = 0.73), followed by the cognitive (R = 0.93, C-index = 0.75) and vitality (R = 0.91, C-index = 0.74) domains.

**Figure 1.**
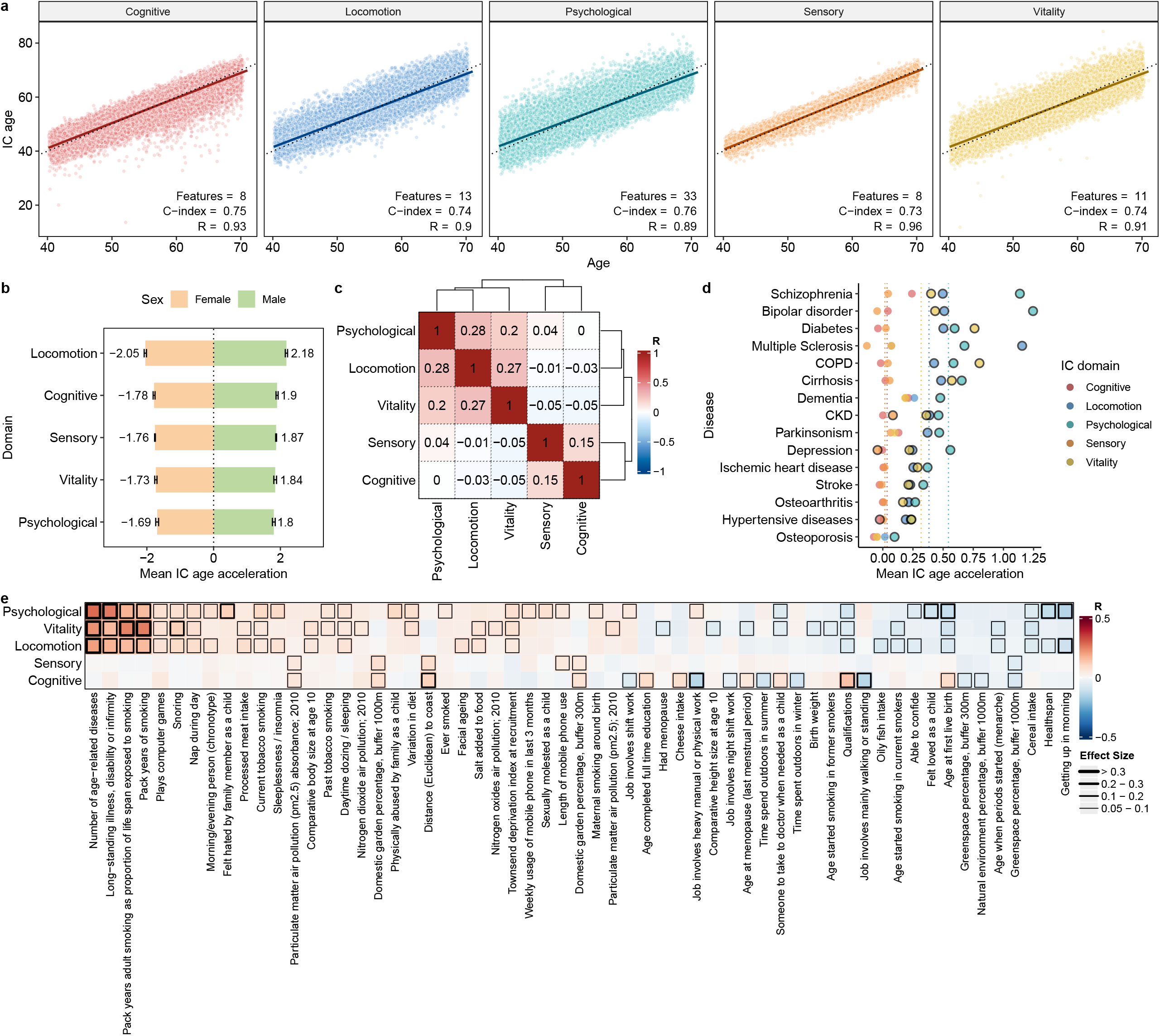
IC age clocks and their associations with diseases and lifestyle. **a**, Relationships between chronological age and predicted IC age across domains of intrinsic capacity. Each plot includes the number of features used (including age and sex), the concordance index (C-index), and the Pearson correlation coefficient (R). **b**, Mean IC age acceleration stratified by sex for each IC domain. **c**, Correlations between IC age acceleration across different IC domains. **d**, Mean IC age acceleration of individuals diagnosed with chronic diseases. Bolded circles indicate age accelerations significantly different from zero. **e**, Correlation between IC domain age acceleration and lifestyle factors. Positive and negative correlations indicate that higher levels of these factors (categorical ordinal or numeric) are associated with increased or decreased IC age acceleration, respectively. Square boxes are displayed with four levels of line width representing increasing effect-size magnitudes.

An analysis of IC age acceleration (IC age adjusted for chronological age) stratified by sex revealed that males exhibit a higher biological age acceleration compared to females across all five domains **(Fig. 1b)**. We found the largest disparity in the locomotion domain, where males show a mean difference in IC age acceleration of 4.23 years compared to females. Across all domains, the psychological domain showed the smallest disparity, though males still exhibited an IC age acceleration increase of 3.49 years compared to females.

Correlations between the IC age acceleration estimates (adjusted for chronological age and sex) revealed two distinct clusters: one composed of the psychological, locomotion, and vitality domains, and the other of the sensory and cognitive domains **(Fig. 1c)**. Notably, the cognitive and psychological domains did not correlate and none of the comparisons showed a positive correlation above 0.28, suggesting that each domain captures different biological aspects in relation to mortality.

The domain-specific IC clocks also display different patterns of association across chronic age-related diseases **(Fig. 1d)**, which validates our hypothesis that domain-specific IC age captures relevant clinical information regarding diseases associated with these domains. For example, the psychological domain shows the highest acceleration in mental health conditions such as bipolar disorder, schizophrenia, and depression. Similarly, vitality age acceleration was associated with COPD, and locomotion age acceleration was associated with multiple sclerosis. Interestingly, most diseases did not show any effect on the acceleration of the cognitive or sensory domains.

We also analyzed the correlations between the IC age acceleration and different lifestyle factors **(Fig. 1e)**. On average, the number of age-related diseases showed the highest correlation with IC age acceleration, particularly affecting the cluster of psychological, vitality, and locomotion domains. Similarly, smoking status (present and past) and pack years were correlated with acceleration in these domains. Feeling hated or harassed as a child, mobile phone usage, and maternal smoking around birth were associated with higher acceleration, particularly in the psychological domain, while we observed the opposite when the participants reported having felt loved as a child. Higher levels of air pollution (nitrogen dioxide, particulate matter) were linked with higher vitality age acceleration, whereas oily fish intake correlated with a decrease in locomotion age acceleration. Lifestyle factors influenced the sensory age acceleration the least; however, we observed a positive correlation with length of mobile phone use and air pollution. Living far from the coast, cheese intake, and qualifications were linked to cognitive age acceleration, while the opposite was observed with the time spent outdoors (summer and winter) and jobs involving physical work. Regarding the chronotype, getting up in the morning was associated with lower acceleration of the psychological and locomotion domains compared to waking in the evening. These associations further highlight the influence of the exposome in potentially shaping intrinsic capacity trajectories across the lifespan.

We also evaluated the differences in IC age acceleration between individuals taking or not taking certain medication regularly (most days of the week for the last 4 weeks) **(Sup Fig. 1)**. While several medications were associated with an increase in age acceleration, most likely reflecting morbidity, a few displayed small but significant decreases in age acceleration. For example, individuals regularly taking vitamin A displayed a decrease of 0.28 years in cognitive IC age. Similarly, regular selenium intake is linked with a decrease in 0.3 years in locomotion IC age. Also, individuals who regularly take glucosamine and fish oil show a decrease between 0.17-0.21 years in vitality and locomotion IC age.

To better understand the molecular mechanisms involved in each IC domain predictor, we performed penalized regression (Lasso regression) on the plasma proteomics data available in the UKB using IC age as the outcome **(Fig. 2a)**. This approach selects the combination of proteins most predictive of IC age in each domain by shrinking non-relevant protein coefficients to zero. Of the 1984 proteins associated with at least one domain, 37% (736 proteins) were linked with just one domain, while 14% (287 proteins) were relevant for all five domains **(Sup Fig. 2a)**. Interestingly, proteins associated with more systems increasingly showed higher correlation with age **(Sup Fig. 2b)**, indicating that common proteins detected in blood across IC age surrogates tend to capture chronological age, while domain-specific proteins tend to capture changes associated with intrinsic capacity domains. By analyzing the domain-specific proteins **(Fig. 2b)**, we linked IGFBPL1, a master regulator of microglia homeostasis^30^, to cognitive IC age, and MYOC (Myocilin), a regulator of muscle and bone formation^31,32^, to locomotion IC age. Similarly, Acid Sphingomyelinase (SMPD1) activity, which relates to major depressive disorder^33^, contributed to psychological IC age, while Cadherin-23 (CDH23), a structural component of the inner hair cells and one of the most studied causal genes in hearing loss^34^, influenced sensory IC age. Finally, NTRK2 (also known as TrkB), a receptor for brain-derived neurotrophic factor (BDNF), and regulator of energy metabolism^35^, was associated with vitality IC age. We also performed pathway enrichment analysis to identify pathways associated with proteins linked with each domain **(Fig. 2c)**. Of the 65 pathways associated with at least one domain of IC, 18 (27%) were relevant for all domains, including canonical signaling pathways such as PI3K-Akt, MAPK, JAK-STAT, and Ras. Among these pathways, the Epidermal Growth Factor Receptor (EGFR) acted as a common regulator.

**Figure 2.**
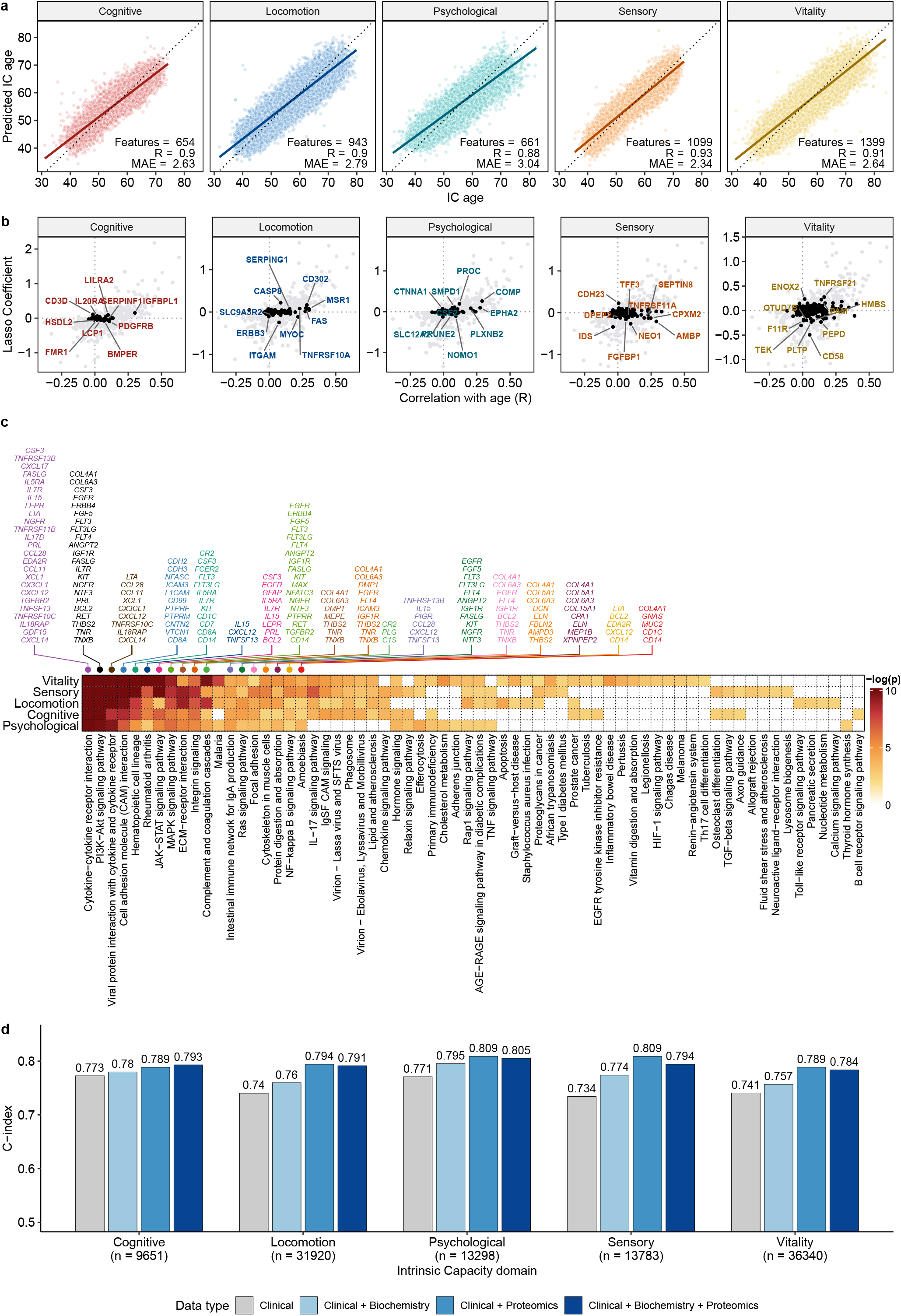
Proteomic surrogates of IC age. **a**, Proteomics-based estimation of IC age across functional domains. Scatter plots show the correlation between chronological age and IC age predicted from proteomics. Each graph includes the number of proteins selected by Lasso regression, the Pearson correlation (R), and the Mean Absolute Error (MAE). Solid lines represent the linear regression fit, and dashed lines indicate the identity line. **b**, Domain-specific proteomic biomarkers and their association with chronological age in each IC domain. **c**, Pathways associated with predictive features in each domain. Heatmap values represent the significance (on a log scale) of a KEGG pathway over-representation test. Top annotations highlight biological pathways relevant to all domains and the shared features across domains. **d**, Comparative performance of mortality prediction models using different types of data aggregated. Sample sizes for each domain are specified on the x-axis.

We evaluated whether combining proteomic and biochemical and hematological markers associated with each IC age domain could improve mortality prediction performance beyond clinical data alone **(Fig. 2d)**. We used a set of proteomic or biochemical features that were predictive of IC age in different domains and merged them with the clinical features to predict mortality risk **(Sup Table. 3)**. We designed the model to keep clinical features fixed in the model, while shrinking biochemical or proteomic features associated with each IC domain if they did not predict mortality. Overall, we observed that adding biochemical features improved the C-index on average by a modest 0.021 while proteomic features increased it by 0.046. However, including both biochemical and proteomic features did not improve the mortality prediction beyond proteomic features alone (Mean C-index = 0.798).

Our study demonstrates that mortality risk estimators built on intrinsic capacity domains capture the specific signal of IC decline through meaningful lifestyle and disease associations. For example, individuals with bipolar disorder, depression and schizophrenia show significantly higher psychological IC ages. This metric also captures associations of psychological IC age with early life exposures such as childhood adversity and maternal smoking around birth. We also demonstrated conserved sex disparities in IC, with males exhibiting significantly higher levels of decline in all five domains compared to females, aligning with previous observations of organ aging where males typically show faster biological aging across most organs^36^. By analyzing proteomics data, we identified known protein biomarkers of IC function such as CDH23 for sensory capacity. These biomarkers may serve as candidates for future mechanistic and causal investigation. We also developed proteomic surrogates of these domain-specific predictors, to enable a quantitative assessment of intrinsic capacity decline through a blood sample. These findings support a shift from disease-centric models toward functional aging frameworks, where decline in intrinsic capacity precedes clinical disease^37^ and reflects heterogeneous deterioration across physiological systems. This study has several limitations. All models were developed within the UKB; thus, replication in a demographically diverse cohort is needed. Also, given that only cross-sectional data is used, the associations with lifestyle factors and medication do not reflect causation, and if they do, could reflect reverse causation. Similarly, proteomic and biochemistry surrogates derived with penalized regression select features to maximize prediction and not to establish causation, thus these should be interpreted as biomarkers instead of mechanistic drivers.

## Methods

### Operationalization of Intrinsic Capacity

The study used data from the UK Biobank (UKB), a large-scale prospective cohort including individuals between 37 - 73 years old during the recruitment phase^38^. Based on a previous systematic review^17^ and a previous IC score calculated in the UKB^39^, we selected features from the UKB across five intrinsic capacity domains (cognitive, vitality, locomotion, sensory and psychological). We only considered features in each domain that were measured in at least 200,000 individuals. In addition to age and sex, we extracted 63 features from the UKB. The cognitive domain integrated six assessments of executive function and memory, including trail-making duration and fluid intelligence scores. The vitality domain comprised nine physiological and metabolic measures, such as spirometry (FEV1, FVC, PEF), grip strength, and biomarkers including hemoglobin concentration and IGF-1. Locomotion was characterized by 11 features including self-reported walking pace, fall history, and metabolic equivalent of task (MET) minutes for various activity intensities. The sensory domain included six features related to visual and hearing aids, auditory performance and eye disorders. Finally, the psychological domain included 31 features capturing mental health, personality traits, and life satisfaction. For each domain, the analyses were restricted to individuals with complete data on all domain-specific features (no imputation was performed).

### Mortality modeling and IC age calculation

To predict mortality using intrinsic capacity domain information, we used Cox Proportional Hazard modelling using the survival^40^ and caret^41^ packages in R. Mortality data (Fields 40000, 40001, and 40007) included 55,023 deceased individuals with follow-up of up to 18.3 years. Model performance was assessed via five-fold cross-validation with folds stratified by age and sex to ensure demographic consistency across iterations. Predictive accuracy for each domain was quantified using Harrell’s concordance index (C-index)^42^ calculated from the estimations in out-of-fold sets. To enhance interpretability of the mortality risk estimates, the values were transformed into age units by matching the distribution of mortality risk and chronological age. This monotonic rescaling does not alter the ranking of individuals and resulted in a transformed value we called “IC age”, which is a mortality-risk estimate expressed in age-like units, and can be interpreted as the biological age of an individual based on their IC domains. IC age acceleration was then calculated for each individual as the residual of a linear regression between the IC age and chronological age adjusting by sex.

### Evaluation of disease-specific intrinsic capacity decline

To calculate the associations between IC age acceleration in different domains and disease prevalence, we integrated three sources of diagnostic data: self-reported medical conditions (Field 20002) and clinical diagnoses recorded via ICD-9 (Field 41271) and ICD-10 (Field 41270) codes. Out-of-fold estimates of IC age were used for the analysis. IC age acceleration values (IC age adjusted by chronological age and sex) were standardized within each domain. We determined if the mean IC age acceleration for each domain and disease pair deviated from zero using one-sample t-tests. To adjust for multiple testing, p-values were adjusted using the Benjamini-Hochberg False Discovery Rate (FDR) method^43^.

### Correlation of lifestyle and medication with domain-specific biological age

To characterize lifestyle profiles, we extracted socioeconomic, environmental and behavioral features from the UKB, while ensuring no overlap with the previously defined features of the intrinsic capacity domains. To ensure comparability across the features, the categorical variables were recoded so that the higher value represented a higher level of the feature described. For example, educational qualifications (Field 6138) were transformed into a six-point ordinal scale where 1 corresponded to CSEs and 6 represented university degrees. For the association analysis, we calculated the Pearson correlation coefficient between the IC age acceleration (sex-adjusted) for each domain and the processed lifestyle features. Given the high statistical power of the UKB, even small effect sizes were statistically significant (FDR <0.05); however, we only displayed those features that exhibited an absolute correlation above 0.05 in at least one intrinsic capacity domain. The analysis of changes in IC age associated with intake of medication was done using field 100045 in the UK Biobank. Individuals who declared none of the above were used as controls. The statistical significance of the difference was calculated using a Wilcoxon test and p-values for all the tests across domains were adjusted by multiple testing correction using the Benjamini & Hochberg method.

### Proteomic drivers of domain-specific IC age

To identify proteins associated with IC age in each domain, we used Lasso regression modeling (alpha = 1) implemented in the glmnet^44^ package in R. We performed 10-fold cross-validation across a range of lambda (10^-5^ to 10^5^) values using mean absolute error as the loss function. To create a more generalizable model, we selected the model with a lambda within one standard error of the minimum error. Proteins associated with each intrinsic capacity domain used for the downstream analyses corresponded to those with non-zero coefficients in the final model. The same approach was used to identify biochemical markers associated with IC age.

### Enrichment analysis of IC-associated proteins

To identify pathways linked with each intrinsic capacity domain, we performed an over-representation test using the clusterProfiler package^45^. First, the gene symbols of proteins with non-zero coefficients in the Lasso models^46^ were converted into Entrez identifiers using the org.Hs.eg.db database^47^. We tested for enriched human pathways within the KEGG database^48^. Pathways were restricted to a gene set size between 10 and 500 and were considered significant if the adjusted p-value was less than 0.05. The resulting p-values were visualized across IC domains using a heatmap generated with the ComplexHeatmap package^49^.

### Integration of molecular and clinical features for mortality prediction

To determine if molecular data improved the mortality prediction, we developed models integrating blood biochemistry and plasma proteomics features with the clinical features of intrinsic capacity. We used Cox penalized regression implemented in the glmnet package^46^. We set penalty factors to zero for clinical features (age, sex and IC domain variables) to ensure they were always retained in the model, while allowing the Lasso regression to perform variable selection on the additional proteomic and biochemical markers. Model performance in predicting mortality was evaluated using Harrell’s Concordance Index (C-index) across four configurations: (1) clinical features only, (2) clinical and biochemistry, (3) clinical and proteomics, and (4) clinical, biochemistry and proteomics data. Predictive accuracy was measured for each domain using samples where all the data were available.

## Data Availability

The UK Biobank data are available under controlled access. Researchers requiring access should apply (https://www.ukbiobank.ac.uk/enable-your-research/apply-for-access) and pay a fee depending on the tier of data requested.

## Code Availability

All custom code used for the analyses is available on GitHub: https://github.com/msfuentealba/ICage.

## Acknowledgements

This work was supported by the National Institutes of Health (NIH) through grants U01 AG086214 (D.F., M.F) and R03 OD036497 (D.F).

## Author Contribution Statement

M.F. and D.F. conceptualized and designed the study. M.F. performed the bioinformatics analyses. D.F. supervised the work. M.F. drafted the manuscript. T.Z, S.A.A.D, V.N.G. and M.P.S. contributed to the conceptual interpretation of the results, provided scientific input and discussion. All authors reviewed and approved the final version.

