## Supplementary Figures for "Quantification of domain-specific intrinsic capacity using mortality data"

Supplementary Figure 1. Age acceleration differences in individuals regularly taking certain medication. Individuals reporting not taking the medication were used as controls. Circled values represent statistically significant differences (Wilcoxon-test) after FDR correction.

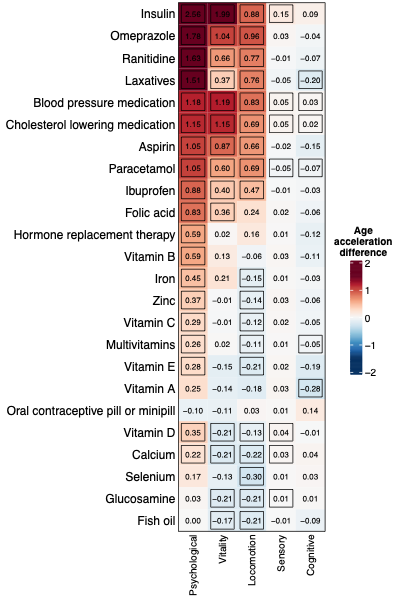

Supplementary Figure 2.a) Number of proteins associated with different sets of domains. b) Absolute Pearson’s correlation of proteins associated with different number of domains.

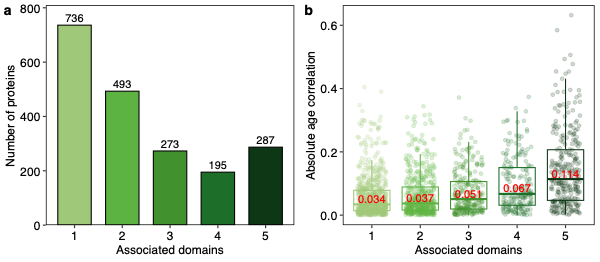
